# Two types of locus coeruleus norepinephrine neurons drive reinforcement learning

**DOI:** 10.1101/2022.12.08.519670

**Authors:** Zhixiao Su, Jeremiah Y. Cohen

## Abstract

The cerebral cortex generates flexible behavior by learning. Reinforcement learning is thought to be driven by error signals in midbrain dopamine neurons. However, they project more densely to basal ganglia than cortex, leaving open the possibility of another source of learning signals for cortex. The locus coeruleus (LC) contains most of the brain’s norepinephrine (NE) neurons and project broadly to cortex. We measured activity from identified mouse LC-NE neurons during a behavioral task requiring ongoing learning from reward prediction errors (RPEs). We found two types of LC-NE neurons: neurons with wide action potentials (type I) were excited by positive RPE and showed an increasing relationship with change of choice likelihood. Neurons with thin action potentials (type II) were excited by lack of reward and showed a decreasing relationship with change of choice likelihood. Silencing LC-NE neurons changed future choices, as predicted from the electrophysiological recordings and a model of how RPEs are used to guide learning. We reveal functional heterogeneity of a neuromodulatory system in the brain and show that NE inputs to cortex act as a quantitative learning signal for flexible behavior.

## Introduction

In dynamic environments, the brain learns from the outcomes of actions. In the framework of reinforcement learning, reward prediction errors (RPEs) guide decisions by updating the internal models used to generate behavioral policies. Experiments across mammals found that midbrain dopamine neurons signal RPE (Schultz et al., 1997; Bayer and Glimcher, 2005; Cohen et al., 2012; Schultz, 2015; Watabe-Uchida et al., 2017). However, there is a puzzle: in rodents, dopamine neurons project densely to the basal ganglia, but much less to cortex (Descarries et al., 1987; Nomura et al., 2014). How, then, does cortex know about the errors used to guide flexible learning? One possibility is that information must propagate through basal ganglia before arriving in cortex. Another is that cortex receives a direct input carrying RPE, distinct from dopamine neuron input to the basal ganglia.

The pontine locus coeruleus (LC) supplies norepinephrine (NE, also known as noradrenaline) to most of the forebrain, including the cerebral cortex (Swanson, 1976; Amaral and Sinnamon, 1977; Jones and Moore, 1977; Foote et al., 1983). In contrast to dopamine neurons, LC-NE neurons project extensively throughout cortex, but very little to basal ganglia (Swanson and Hartman, 1975; Nomura et al., 2014). LC-NE neurons are thought to modulate arousal, attention, exploratory decision making, and network activity in cortex (Aston-Jones et al., 1997; Berridge and Waterhouse, 2003; Bouret and Sara, 2004; Aston-Jones and Cohen, 2005; Bouret and Sara, 2005; Dayan and Yu, 2006; Sara, 2009; Kalwani et al., 2014; Tervo et al., 2014; Bouret and Richmond, 2015; Bornert and Bouret, 2021).

Several observations point to heterogeneity of LC-NE neurons. LC neurons with different somatic morphologies show biases in their axonal projections (Loughlin et al., 1986). *Ex vivo* (Williams et al., 1984; Chandler et al., 2014) and *in vivo* (Totah et al., 2018) electrophysiological recordings show different biophysical features in LC neurons. LC-NE neurons with different projection targets receive different inputs (Schwarz et al., 2015) and have different functions on target areas (Hirschberg et al., 2017; Uematsu et al., 2017). Together, these observations indicate potentially important functional heterogeneity among LC-NE neurons.

Here, we show that two subtypes of LC-NE neurons modulate reinforcement learning in distinct ways. We measured and manipulated LC-NE activity in mice performing a task that required adapting behavior as probabilities of reward given alternative actions changed. One subtype provides a quantitative RPE for frontal cortex to learn in dynamic environments.

### Mice learn from recent experience to make future choices

We trained head-restrained mice on a dynamic decision-making task (Figures 1a and A1; Grossman et al., 2022). Each day, mice performed a few hundred trials (range, 153–602 trials; Figure A1c). Each trial began with a 500-ms tone “go cue” (or, in 5% of trials, a control “no go” cue). On each trial, mice chose freely between licking toward a leftward or rightward tube. Each tube probabilistically delivered a water reward (*P*(*R*)) following a 300-ms delay after choice. The outcome (presence or absence of reward) was followed by a variable inter-trial interval before the onset of the next trial.

**Figure 1.**
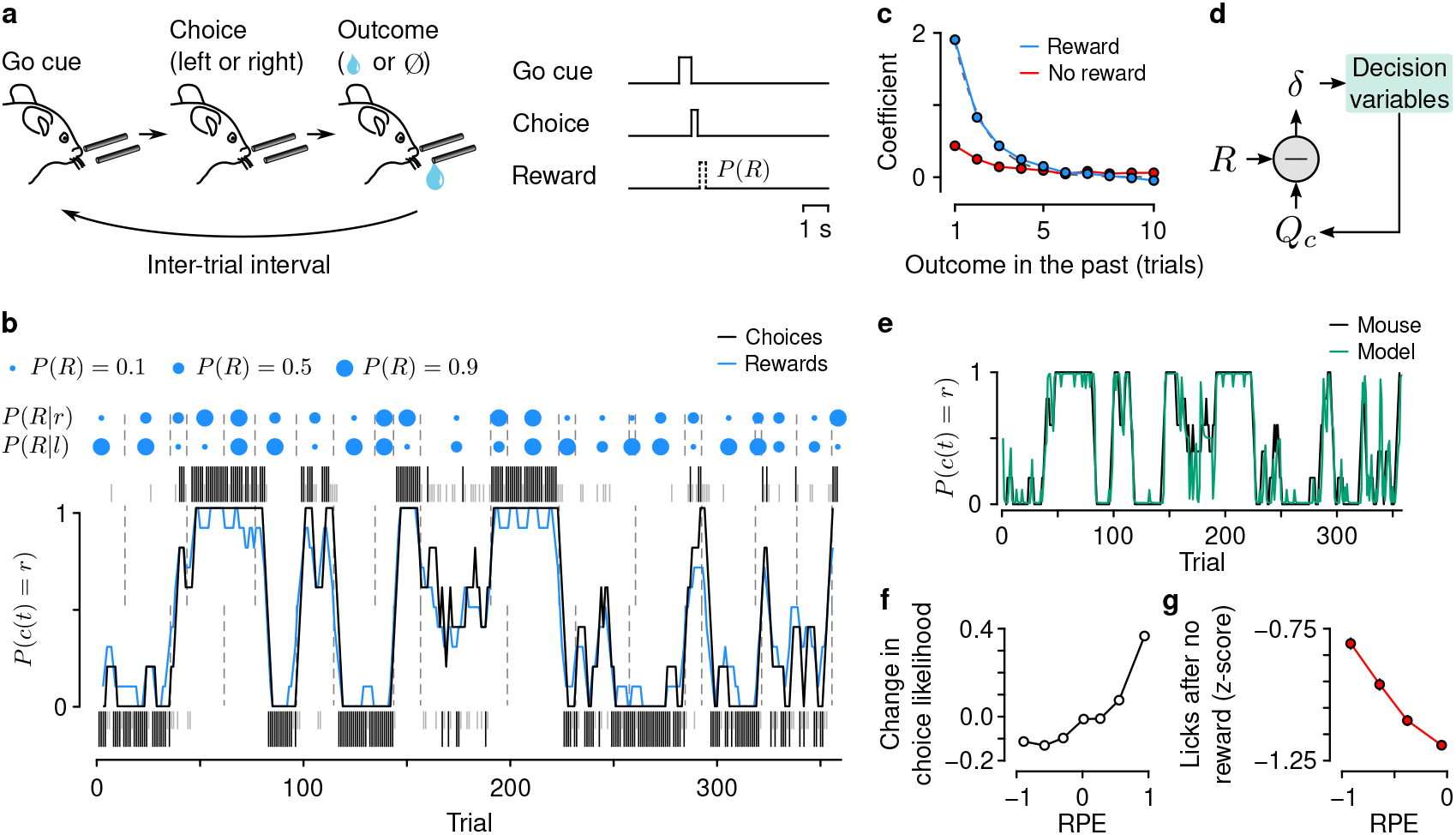
Mice forage dynamically for rewards. (a) Dynamic foraging task in which mice chose freely between a leftward and rightward lick after a “go cue,” followed by a reward with a probability (*P*(*R*)) that varied over time. (b) Example mouse behavior from a single session in the task. Tall black (rewarded) and short gray (unrewarded) ticks correspond to left (below) and right (above) choices. Black curve: mouse (smoothed over 5 trials, boxcar filter) choices. Blue curve: rewards (smoothed over 5 trials). Blue dots indicate left/right reward probabilities (*P*(*R*|*l*) and *P*(*R*|*r*), respectively) and dashed lines indicate a change in *P*(*R*). (c) Logistic regression coefficients for choice as a function of outcome history for all mice. Error bars: 95% CI. *R*^2^ = 0.60, successful choice prediction, 0.88. Gray: exponential fit (coefficient of 3.95). (d) Reinforcement-learning model. The predicted value of a choice (*Q_c_*) is compared to reward (*R*) to generate a *reward prediction error* (*δ*, RPE). (e) *Q*-learning fit to example mouse behavior from (b). Success rate of predicting choices, 0.87. Per-trial log likelihood, –0.24. (f) Relationship between the change of likelihood of choice as a function of RPE (rank correlation, *ρ* = 0.84, *p* = 0.0). Change of choice likelihood is defined as 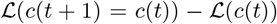, where 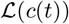 is the likelihood of choice on trial *t* and 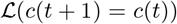 is the likelihood of *c*(*t*) on trial *t* + 1. This quantity measures how choice dynamics change on consecutive trials. (g) Mice showed higher lick rates following no reward with increasing negative RPE (mean ± SEM, 23 mice, 652 sessions; linear regression coefficient, –0.54 ± 0.019).

Reward probabilities changed in blocks of trials for leftward and rightward choices, inducing mice to learn from experience. In this choice task, mice behaved dynamically, tracking the changing *P*(*R*) with their choice patterns (Figure 1b). Mice used reward and choice history to influence future choices (Figure 1c). We quantified learning using *Q*-learning, a reinforcement-learning model that estimates the values of alternative actions and learns from errors in expected action outcomes (RPE; Figure 1d; Bertsekas and Tsitsiklis, 1996; Sutton and Barto, 1998). The model provides an estimate of RPE on single trials (Figures 1e and A1).

Interestingly, mice appeared to use RPE to update choice strategy on the next trial as well as modify ongoing movements. There was a monotonically increasing relationship between the change of choice likelihood as a function of RPE (Figure 1f), indicating that mice used RPE from one trial to influence choice on the next. Upon receiving no reward, mice also showed higher lick rates when RPE was more negative (that is, when they did not receive reward but strongly expected it), suggesting that RPE was also used to modify ongoing movements (Figures 1g and A1f). With a quantification of learning in hand, we turn our attention to LC-NE neurons.

### Two types of LC-NE neurons

To measure the activity of LC-NE neurons, we made electrophysiological recordings from LC of mice expressing Cre recombinase under the control of dopamine beta hydroxylase (*Dbh*-Cre), the rate-limiting enzyme for NE production (Figures 2a and 2b). We expressed the light-gated ion channel, channelrhodopsin-2 (ChR2), selectively in NE neurons using injections of adeno-associated virus under the control of Cre recombinase (AAV-DIO-ChR2-EYFP). Together with microelectrodes, we implanted optic fibers to deliver 473-nm light to the region surrounding the electrodes. We identified LC-NE neurons as those with shortlatency and high-probability responses to light (Figures 2c, 2d, and A2). Using these criteria, we recorded from 149 LC-NE neurons from 4 mice during the behavioral task.

**Figure 2.**
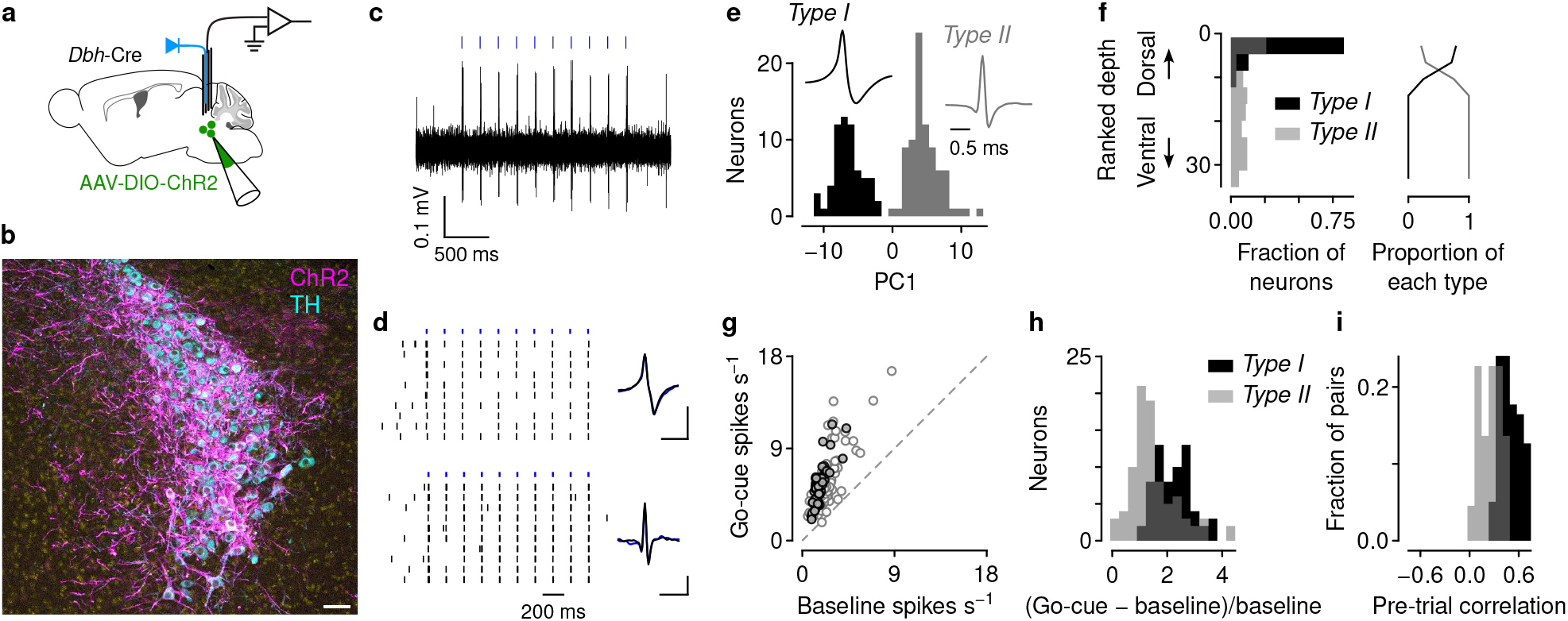
Two types of LC-NE neurons. (a) Schematic of electrophysiological recordings from identified LC-NE neurons in *Dbh-Cre* mice. (b) Expression of ChR2 colocalizes with LC-NE neurons (identified by tyrosine hydroxylase [TH] immunostaining). Scale bar: 50 *μ*m. (c) Example train of action potentials in an LC-NE neuron evoked by a sequence of light pulses. (d) Example responses of two LC-NE neurons to repeated trains of 20-ms light stimuli (across rows). Inset: spontaneous (black) and light-evoked (blue) action potential waveforms. Scale bars: 0.1 mV, 1 ms. (e) PCA on action potential waveforms yields two distinct clusters: neurons with wide (type I, 63 neurons) or thin action potentials (type II, 86 neurons). (f) Type I neurons were found relatively more dorsal within LC than type II neurons (Wilcoxon rank sum test, *p* = 3.6 × 10^−18^). Ranked depth was estimated within-mouse and aggregated across mice. (g) All LC-NE neurons are excited following go cues relative to baseline (paired t-test, *t*_148_ = 23.9, *p* = 1.1 × 10^−52^). (h) Change in go-cue response relative to pre-trial baseline firing rates, normalized by baseline. Type I neurons are more excitable than type II neurons (t-test, *t*_147_ = 5.59, *p* = 1.1 × 10^−7^). (i) Activity of pairs of simultaneously recorded type I neurons were more correlated than pairs of type II neurons (t-test, *t*_99_ = 8.12, *p* = 1.3 × 10^−12^).

We observed two populations of LC-NE neurons with distinct properties. One group of neurons (“type I”) had wide action potentials, whereas the other had thin action potentials (“type II”; Figures 2e and A2a). Consistent with studies in rats, we found that type I neurons were more dorsal in LC than type II neurons (Figure 2f; Totah et al., 2018). Although all LC-NE neurons showed a brief excitation in response to go cues (Aston-Jones et al., 1994; Bouret and Sara, 2004; Clayton et al., 2004; Yang et al., 2021), type I neurons showed comparatively larger go-cue-evoked responses than type II neurons (Figures 2g and 2h). Type I neurons also showed higher correlations in firing rates among simultaneously recorded pairs than type II neurons (Figures 2i and A2d). The latter observation agrees with recent *in vivo* (Totah et al., 2018) and *ex vivo* (McKinney et al., 2022) experiments, indicating that LC-NE neurons are embedded in subtype-specific networks.

### Subtypes of LC-NE neurons are either excited by RPE or lack of reward

Do type I and type II neurons behave differently as mice learn from action outcomes? Figure 3a shows an example of each subtype of LC-NE neuron. Following the choice lick, the type I neuron showed a monotonic increase in firing rate with increasing RPE. By contrast, the type II neuron was excited by lack of reward, without modulation by expected reward (that is, *Q_c_*). Importantly, these response dynamics were evoked without any external stimulus, other than the presence or absence of reward.

**Figure 3.**
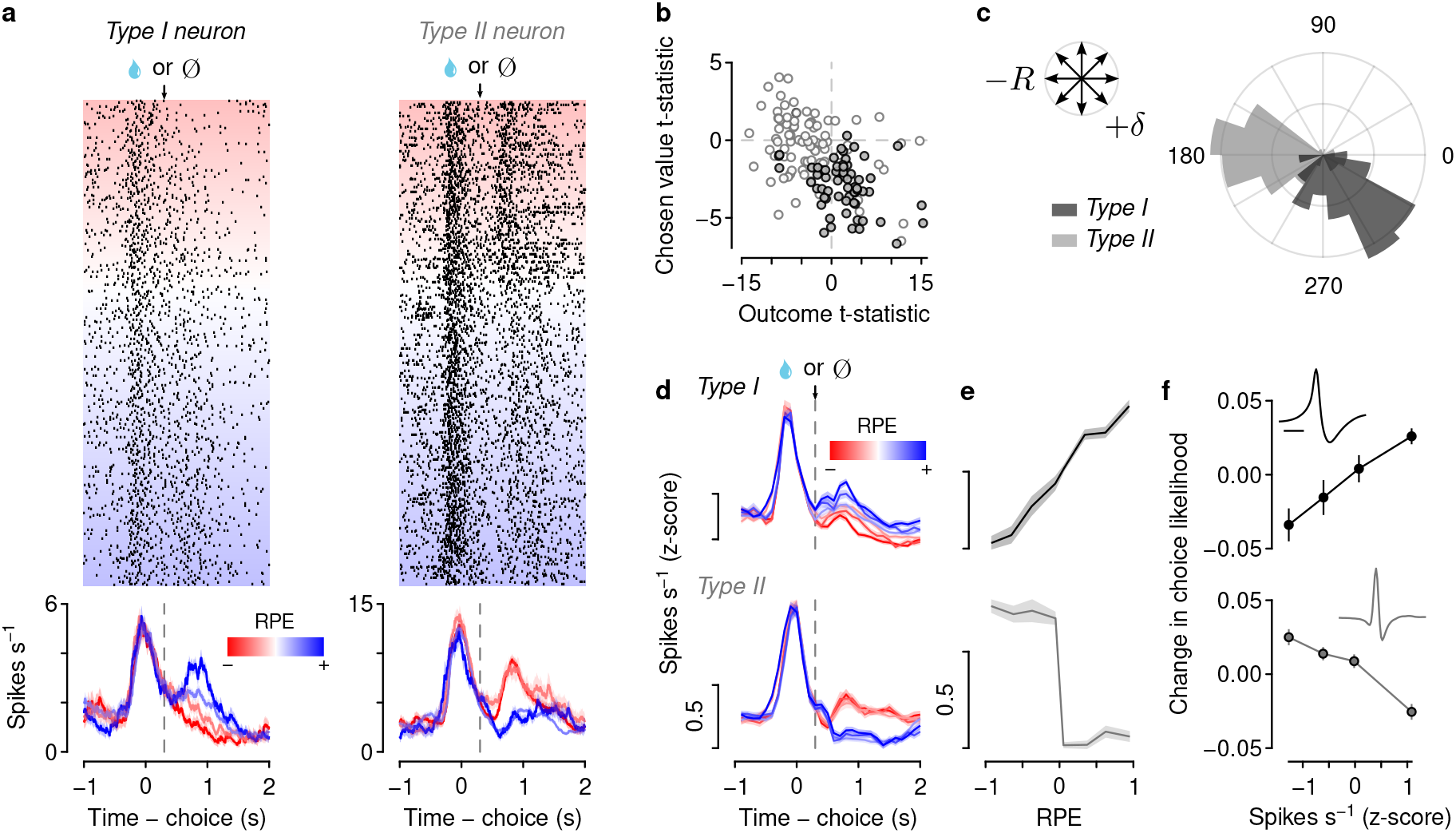
Type I LC-NE neurons scale with RPE whereas type II LC-NE neurons are excited by lack of reward. (a) Example type I (left) and type II (right) neurons. Upper plots show action potentials (ticks) for each trial (rows), sorted by RPE. Lower plots show mean ± SEM firing rates in RPE quartiles. Note that the type I neuron’s response scales with RPE, whereas the type II neuron is excited by lack of reward regardless of chosen value. (b) Effects of chosen value versus outcome (reward or no reward) on firing rates of type I (black) and type II (gray) neurons. Each t-statistic comes from a regression of a neuron’s firing rates on chosen value and outcome. (c) Polar histogram of coefficients from regression models in (b) shows different dynamics among LC-NE subtypes: type I neurons correlate with RPE (lower right portion of the polar plot, +*δ*), whereas type II neurons correlated with no reward (left portion of the polar plot, –*R*). Concentric circles correspond to 0.1 density. Kuiper’s test, *k* = 4.0 × 10^3^, *p* = 10^−3^. (d) Mean z-scored firing rates of type I (28 neurons) and type II (64 neurons) LC-NE neurons with significant regression coefficients from (b). (e) Activity of each LC-NE subtype versus RPE. Type I neurons scale linearly with RPE, whereas type II neurons behave more like a step function: excitation to lack of reward. (f) Higher activity of type I neurons correlated with larger change in choice likelihood 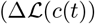; linear regression coefficient, 0.032; *t* = 13.5, *p* = 1.2 × 10^−26^). Type II neurons showed the opposite effect (coefficient, –0.020; *t* = 11.0, *p* = 4.6 × 10^−23^). Insets reproduce mean action potential waveforms from Figure 2e.

Across the population, types I and II LC-NE neurons showed distinct responses during behavior: RPE or lack of reward (see Figure A3 for additional examples). We calculated regressions of firing rates during the outcome (reward or no reward) as a function of the expected reward value of the choice (*Q_c_*) and outcome (Figure 3b). Type I and type II neurons clustered in different regions of this plot: type I neurons were primarily excited by reward and inhibited by *Q_c_* (that is, they correlated with RPE), whereas type II neurons were primarily excited by lack of reward without strong modulation by *Q_c_*. To quantify this observation, we calculated the angle of the coefficients from the regressions in Figure 3b relative to the origin. The resulting polar histogram revealed distinct clustering of the task-related activity of type I and type II LC-NE neurons (Figure 3c). Average responses of neurons of each type showed modulation by RPE or lack of reward (Figures 3d and 3e).

### LC-NE neuron silencing changes future choices, consistent with a change in RPE

Are RPEs in LC-NE neurons used to update future choice behavior? If so, their activity should correlate with future behavior and silencing LC-NE neurons during a trial should change choice behavior on future trials. We first calculated the change in choice likelihood 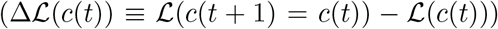 and compared it against the activity of LC-NE neurons. We counted action potentials during the outcome, when mice experienced RPE. We found two distinct profiles in the two types of LC-NE neurons. Type I neurons showed an increasing relationship between 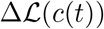 and firing rates: higher firing rates predicted positive changes of choice likelihood (Figure 3f). Type II neurons showed the opposite relationship: a monotonic decrease of 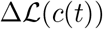 as firing rates increased.

We next tested whether bilateral silencing of LC-NE neurons disrupted future choices. We expressed the light-gated chloride channel GtACR2 selectively in LC-NE neurons by injecting AAV-SIO-GtACR2-FusionRed into the LC of *Dbh*-Cre mice (Figure 4a). We delivered 473-nm light bilaterally on 30% of trials (Figure 4b). Light delivery effectively decreased the pupil dilation evoked during a trial (Figures 4c and 4d), indicating effective silencing. We used low irradiance (0.8–1 mW) to target more dorsal LC. We reasoned that this would preferentially silence type I versus type II neurons due to their dorsal-ventral distribution (Figure 2f). That is, we wished to preferentially silence RPE arising from LC-NE neurons.

**Figure 4.**
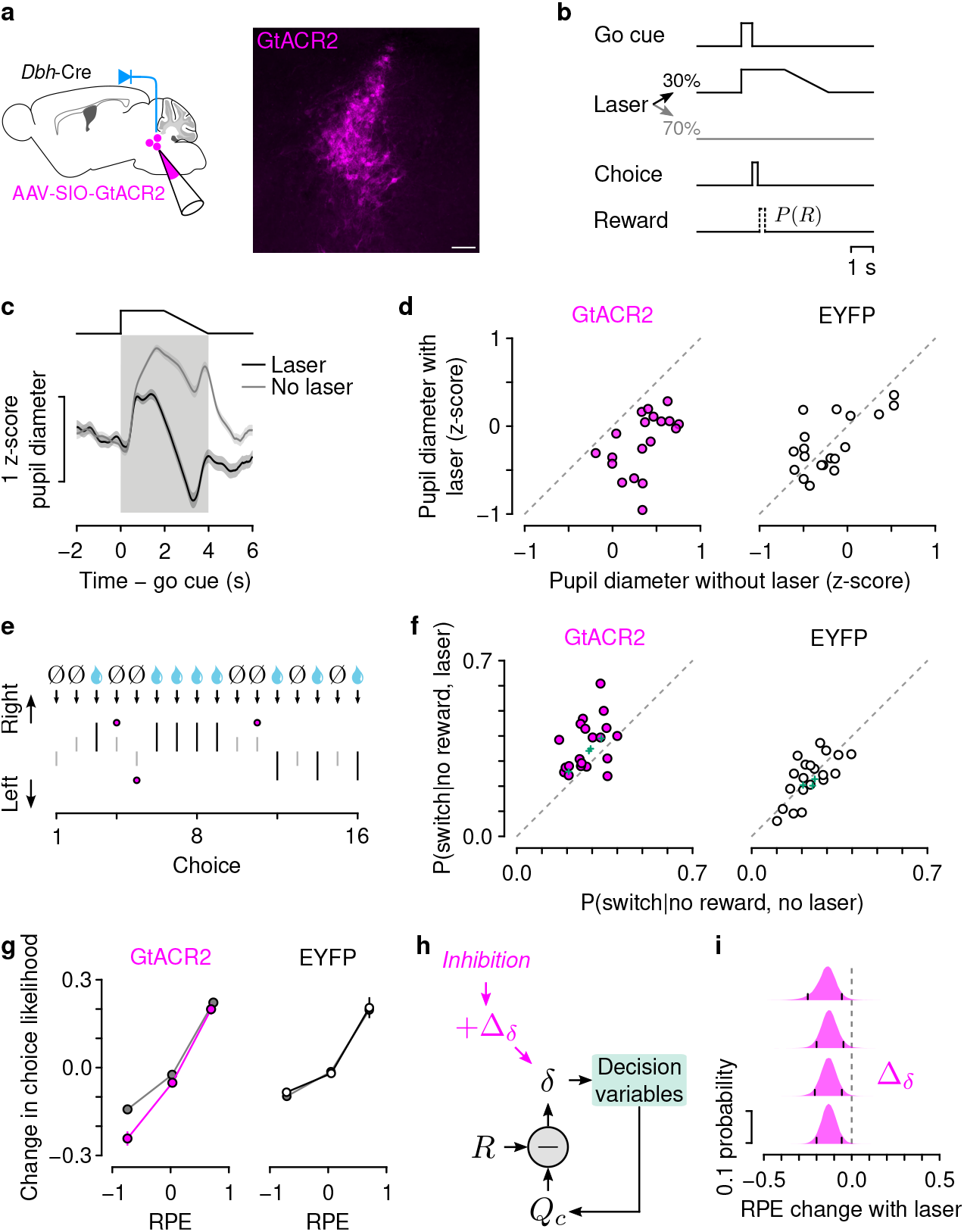
LC-NE silencing mimics a decrease in RPE. (a) Left: strategy to silence LC-NE neurons using GtACR2 expression in *Dbh*-Cre mice. Right: GtACR2 expression in LC-NE neurons (scale bar: 100 *μ*m). (b) Experimental design. Light onset was coincident with go cues on 30% of trials, remained constant for 2 s, and ramped down for 2 s to minimize post-inhibitory rebound excitation. (c) Example experiment with attenuated pupil dilation with LC-NE silencing. (d) LC-NE inhibition attenuated pupil dilation (paired t-tests, *t*_19_ = 7.79, *p* = 2.5 × 10^−7^ for GtACR2; *t*_19_ = 0.26, *p* = 0.80 for EYFP). (e) Example sequence of trials showing switching following no reward with laser versus without. (f) Increased probability of switching after no reward (*P*(switch|no reward)) with LC-NE silencing but not in control experiments. Green: model predictions of *P*(switch|no reward) for each mouse. Wilcoxon signed rank tests: GtACR2, *p* = 0.0029; EYFP, *p* = 0.94. (g) LC-NE silencing caused a negative shift in the change in choice likelihood specifically during negative RPE in mice with GtACR2 (no laser, gray; laser, magenta; signed-rank tests, *p* = 0.00089, *p* = 0.033, *p* = 0.31) but not EYFP (no laser, gray; laser, black; signed-rank tests, *p* = 0.50, *p* = 0.79, *p* = 0.68). (h) Model to estimate the change in RPE (Δ_*δ*_) with LC-NE inhibition. (i) The posterior distribution of Δ_*δ*_ was less than zero in each mouse. Black ticks denote 95% certainty intervals.

Preferential silencing of type I neurons makes two predictions. First, because these neurons signal RPE (Figure 3e), artificially reducing RPE would decrease the value of the chosen action, making it less likely to be chosen in the future. That is, P(switch) should increase. In this task, learning from positive and negative RPEs is asymmetric (Figure A1g; Bari et al., 2019; Grossman et al., 2022). We thus calculated *P*(switch|no reward) and *P*(switch]reward) separately. We found that silencing LC-NE neurons during trials with no reward increased the probability of switching choices (Figures 4e and 4f). In control experiments, we found no change in *P*(switch|no reward) when we expressed a fluorophore alone in LC-NE neurons (Figure 4f). We found no difference in *P*(switch]reward) with LC-NE silencing (Figure A4a). This may be due to strong choice autocorrelation, reflecting a tendency for mice to repeat choices, independent of RPE-driven learning.

The second prediction is that silencing LC-NE RPEs should produce a negative shift in the change of choice likelihood 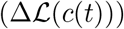, based on the relationship between 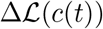 and firing rates (Figure 3f). We indeed found that LC-NE silencing caused a negative shift in 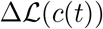 as a function of RPE (Figure 4g). This effect was present only for negative RPE, consistent with the observation that LC-NE silencing increased choice switching following lack of reward (Figure 4f).

To further test whether LC-NE silencing quantitatively mimicked a decrease in RPE, we developed a model in which RPE changed by an additive factor with silencing (Δ_*δ*_ in Figure 4h). For each mouse, the posterior distribution of Δ_*δ*_ was significantly negative, demonstrating that LC-NE silencing mimicked a decrease in RPE (Figure 4i). Importantly, this model predicts that artificially subtracting from RPEs increases *P*(switch|no reward) but not *P*(switch|reward) (green points in Figures 4f and A4a).

We attribute the effects of silencing to reductions in RPE as opposed to a general effect on choice variability or arousal. If LC-NE inhibition simply increased future choice variability, then *P*(stay|reward) should also change. As noted above, we observed no change in *P*(stay|reward) with LC-NE inhibition (Figure A4a). LC-NE inhibition also did not alter response times to go cues (Figure A4b), nor did it change the number of ongoing licks following rewarded or unrewarded choices (Figures A4c and A4d). We also compared the results of the model in which silencing shifts RPE (Figures 4h, 4i, and A4f) to one in which silencing made choices more random (Figure A4g). This model did not indicate a strong change in choice stochasticity with LC-NE inhibition (Figure A4h). Given these observations, we conclude that LC-NE silencing selectively disrupted error-driven learning, while sparing other functions involved in task performance.

### LC-NE axons in frontal cortex signal RPE

Are LC-NE RPEs transmitted to cortex? We answered this question by measuring the calcium dynamics of LC-NE axons in medial frontal cortex. We focused on prelimbic cortex (PrL), in which we previously found persistent firing rates carrying relative choice value in the present behavioral task (Bari et al., 2019). If RPE is used to update decision variables in this task, we expect to observe LC-NE RPE input to PrL. We expressed the calcium sensor GCaMP6s using an axon-targeted viral construct (AAV-DIO-axon-GCaMP6s) in *Dbh*-Cre mice (Figures 5a and A5). We recorded fluorescence dynamics simultaneously in LC and PrL (Figure 5b) by implanting optic fibers above each structure. In LC, we predicted that we would sample primarily from more dorsal LC-NE (i.e., type I) neurons. We estimated the transformation from measured spiking rates (Figure 3e) to GCaMP6s dynamics using a previous estimate of transfer functions in cortex (Figure 5c; Wei et al., 2020). Although this function was derived for somatic calcium dynamics in cortex, it gave us a prediction about the qualitative shape of the dynamics we would expect to see if LC-NE axons in PrL came from type I neurons. In both structures, LC-NE dynamics 1.5 s after choice outcome correlated with RPE (Figures 5d–5f). We quantified this observation in PrL by calculating regressions of axonal calcium dynamics as a function of chosen value and outcome. Regression t-statistics (Figure 5g) and coefficients (Figure 5h) were consistent with modulation by RPE (compare to Figures 3b and 3c). We thus conclude that frontal cortex receives RPE input from LC-NE neurons, likely from type I neurons with somata in dorsal LC.

**Figure 5.**
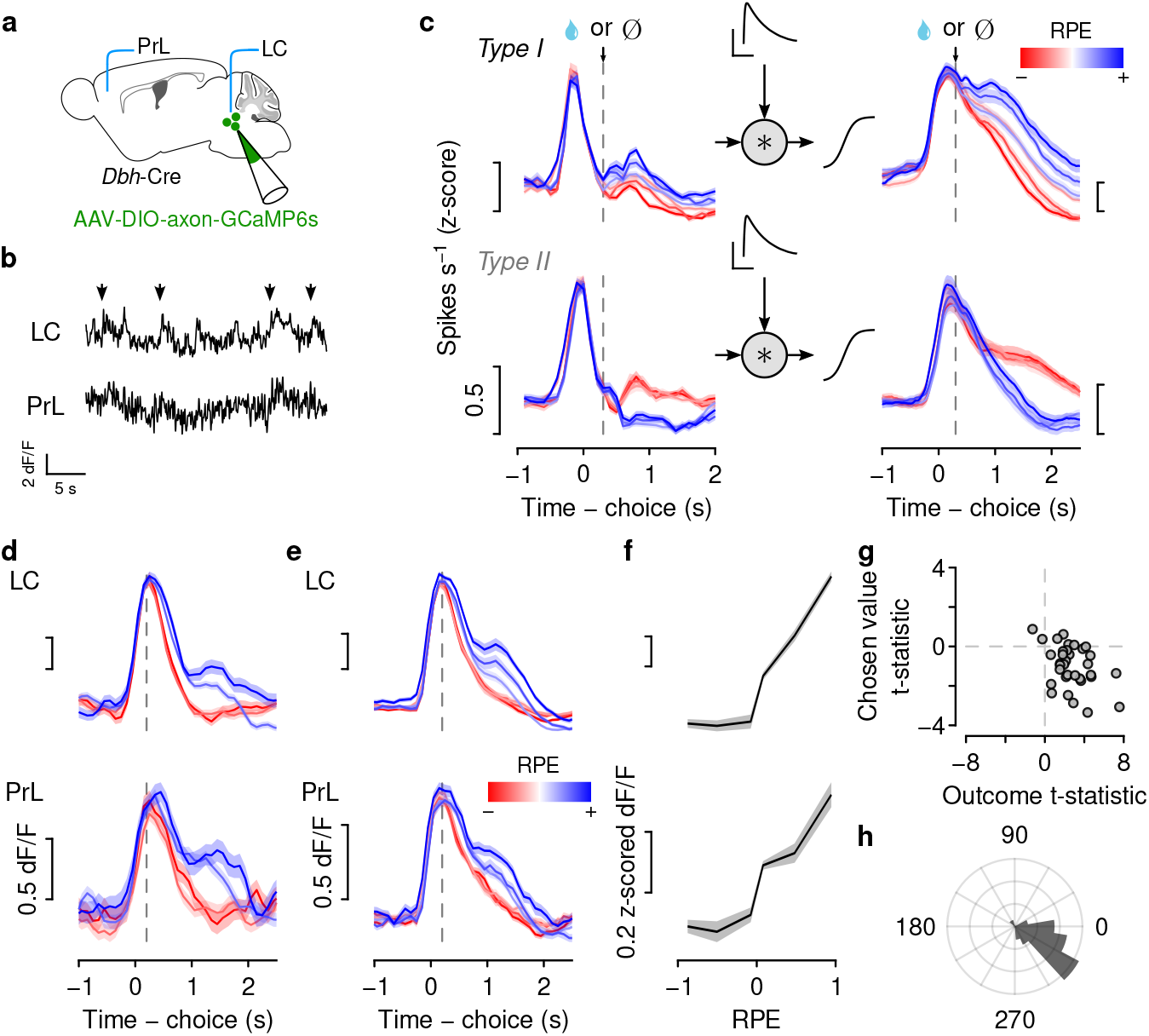
LC-NE dynamics in medial frontal cortex correlate with RPE. (a) Strategy to measure LC-NE calcium dynamics simultaneously in LC somata and their axons in frontal cortex. (b) Example fluorescence in LC and frontal cortex. Arrowheads indicate go cues. (c) Convolving mean type I and type II neuron firing rates (reproduced from Figure 3d) with a kernel (scale bars: 0.5 density, 1 s) and fed into an activation function yields estimated calcium dynamics from population firing rates. (d) Example LC-NE dynamics from one session shows correlations with RPE that are consistent with type I neuron dynamics. (e) Population LC-NE dynamics (2 mice, 34 sessions, mean ± SEM) increase with RPE in both areas. (f) Calcium dynamics increase monotonically with RPE (note that they are rectified relative to spiking activity in Figure 3e, likely due to differences in measurement technique). (g) Effects of chosen value versus outcome on axonal calcium dynamics in PrL (as in Figure 3b). (h) Polar histogram of regression coefficients of axonal calcium dynamics in PrL as a function of chosen value and outcome (as in Figure 3c). Concentric circles correspond to 0.1 density.

## Discussion

We found two biophysically distinct types of LC-NE neurons that had different functions: wide-spiking (type I) neurons signaled RPE, whereas thin-spiking (type II) neurons were excited by lack of reward. Calcium dynamics of LC-NE axons in frontal cortex correlated with RPE. Our silencing experiments are consistent with NE providing a *quantitative* RPE to cortex, a key signal used to drive reinforcement learning.

How does NE input to cortex drive learning? The mechanisms likely include multiple cell types and receptors. NE modulates synaptic plasticity across brain areas (Delaney et al., 2007; Seol et al., 2007; Carey and Regehr, 2009; Hong et al., 2022). *Ex vivo* electrophysiology indicates different effects of NE on different cell types in frontal cortex (Kawaguchi and Shindou, 1998; Dembrow et al., 2010; Dembrow and Johnston, 2014). NE receptors are expressed nonuniformly across cortical cell types (Scheinin et al., 1994; Rosin et al., 1996; Day et al., 1997; Talley et al., 1996; Santana et al., 2013; Santana and Artigas, 2017). In addition, NE may exert its effects on both neuronal and non-neuronal cells (Paukert et al., 2014; Mu et al., 2019), providing a rich substrate for broadcasting an ascending learning signal across large volumes of cortex. This learning signal may generalize beyond reinforcement learning; artificial excitation of LC changes sensory-driven learning as well (Glennon et al., 2019; Jordan and Keller, 2022; McBurney-Lin et al., 2022).

How do synaptic inputs to LC-NE drive RPE computation in type I neurons and excitation following no reward in type II neurons? Major LC afferents include frontal cortex, central amygdala, cerebellum, hypothalamus, and ventral medulla (Cedarbaum and Aghajanian, 1978; Aston-Jones et al., 1986; Luppi et al., 1995; Schwarz et al., 2015). In anesthetized rats, input from frontal cortex (Jodo et al., 1998) and central amygdala (Bouret et al., 2003) excites LC. Excitatory inputs from the medulla (Ennis and Aston-Jones, 1988) may be involved in increased LC firing rates following aversive stimuli (Chiang and Aston-Jones, 1993). It will be important to determine whether biased inputs to subtypes of LC-NE neurons are involved in the responses we observed. Alternatively, afferents to LC-NE neurons may be mixed, and intra-LC networks (Totah et al., 2018; Breton-Provencher and Sur, 2019; McKinney et al., 2022) may produce the distinct outputs we measured here (that is, RPE and excitation to lack of reward).

There is growing knowledge of heterogeneity among LC-NE neurons (Schwarz et al., 2015; Uematsu et al., 2017; Chandler et al., 2019; Poe et al., 2020; Breton-Provencher et al., 2022; Luskin et al., 2022). It remains to be determined whether our two discrete subtypes of LC-NE neurons correspond to known morphological and molecular differences (Swanson, 1976; Loughlin et al., 1986; Luskin et al., 2022; McKinney et al., 2022) or differences in how LC affects different targets (Hirschberg et al., 2017; Uematsu et al., 2017; Breton-Provencher et al., 2022). Mapping structure and function in subtypes of LC-NE neurons will help refine theories of NE in the nervous system, including ideas about exploration, attention, and uncertainty (Aston-Jones and Cohen, 2005; Bouret and Sara, 2005; Dayan and Yu, 2006; Sara, 2009).

We conclude that a major neuromodulatory input to cortex signals RPE, a key variable for reinforcement learning. It is striking that dopamine inputs to basal ganglia carry a similar RPE, yet project comparatively little to cortex (for differences between rodents and primates, see Berger et al., 1991; Lewis et al., 2001). We propose that cortex and basal ganglia may receive complementary inputs from the two major catecholamines in the brain: dopamine and norepinephrine. These two neuromodulators may allow cortex and basal ganglia to operate as hierarchical learning systems (Wang et al., 2018), allowing the brain to learn over multiple timescales (Murray and Escola, 2020).

## Acknowledgments

We thank Drs. Bilal Bari, Daeyeol Lee, and Karel Svoboda for input. This work was supported by R01NS104834 (J.Y.C.) and P30NS050274.

## Author contributions

Z.S. collected data. Z.S. and J.Y.C. designed experiments, analyzed data, and wrote the paper.

## Methods

### Animals and surgery

We used 37 mice, backcrossed with C57BL/6J and heterozygous for Cre recombinase under the control of the dopamine beta hydroxylase gene (Dbh^*tm*3.2(*cre*)*Pjen*^, The Jackson Laboratory, 033951; Tillage et al., 2020). Twenty-three male mice were used for behavioral experiments, five male mice were used for electrophysiological recordings during the dynamic foraging task, seven male mice were used for silencing and control experiments, and two male mice for GCaMP6s recordings. Surgery was performed on mice between the ages of 10–16 weeks, under isoflurane anesthesia (1.5–2.0% in O_2_) and under aseptic conditions. During all surgeries, custom-made titanium headplates were surgically attached to the skull using dental adhesive (C&B-Metabond, Parkell). After the surgeries, analgesia (ketoprofen, 5 mg kg^−1^ and buprenorphine, 0.05–0.1 mg kg^−1^) was administered to minimize pain and aid recovery.

For electrophysiological experiments, we implanted a custom microdrive targeting right LC, entering through a craniotomy at 0.28–0.35 mm anterior to the junction of the inferior colliculus and cerebellum and 0.85–0.90 mm lateral from the midline (identified using the vasculature as a landmark).

For all experiments, mice were given at least one week to recover prior to water restriction. During water restriction, mice had free access to food and were monitored daily in order to maintain 80% of their baseline body weight. All mice were housed in reverse light cycle (12h dark/12h light, dark from 08:00– 20:00) and all experiments were conducted during the dark cycle between 12:00 and 20:00. All surgical and experimental procedures were in accordance with the *National Institutes of Health Guide for the Care and Use of Laboratory Animals* and approved by the Johns Hopkins University Animal Care and Use Committee.

### Behavioral task

Before training on the tasks, water-restricted mice were habituated to head fixation for 1–3 d with free access to water from the two spouts (21 ga stainless steel tubes separated by 4 mm) placed in front of the 38.1-mm diameter acrylic tube in which the mice rested. The spouts were mounted on a micromanipulator (DT12XYZ, Thorlabs) with a custom digital rotary encoder to measure the position of the lick spouts with 5–10 *μ*m resolution (Bari et al., 2019). Each spout was attached to a solenoid (ROB-11015, Sparkfun) to enable movement in the anterior-posterior axis of the mouse. The tones used for the cues (randomly assigned to go and no-go cues per mouse) were 7.5 and 15 kHz square waves generated by microcontrollers (ATmega16U2 or ATmega328), amplified and delivered through speakers (CUI Devices, GF0401M).

Licks were detected using a custom circuit (Janelia Research Campus 2019-053). Task events were controlled and recorded using custom code (Arduino) written for a microcontroller (ATmega16U2 or ATmega328). Water rewards were 2–3 *μ*l, adjusted for each mouse to maximize the number of trials completed per session and to keep sessions around 60 minutes. Solenoids (LHDA1233115H, The Lee Co) were calibrated to release the desired volume of water and were mounted on the outside of the sound-attenuated chamber used for behavior tasks. White noise (2–60 kHz, Sweetwater Lynx L22 sound card, Rotel RB-930AX two-channel power amplifier, and Pettersson L60 Ultrasound Speaker), was played inside the chamber to mask ambient noise.

During the 1–3 days of habituation, mice were trained to lick both spouts to receive water. Water was delivered following a lick to the correct spout at any time. Reward probabilities were chosen from the set {0,1} and reversed every 20 trials.

In the second stage of training (5–12 d), the trial structure with tone presentation was introduced. Each trial began with the 0.5 s delivery of either an auditory “go cue” (*P* = 0.95) or a “no-go cue” (*P* = 0.05). Following the go cue, mice could lick either the left or the right spout. If a lick was made during a 1.8 s response window, reward was delivered probabilistically from the chosen spout. The unchosen spout was retracted when the tongue contacted the chosen spout to prevent mice from sampling both spouts within a trial. The unchosen spout was replaced 3.1 s after cue onset (or 3.0 s in experiments with LC-NE silencing or axonal calcium dynamics measurements). Following a no-go cue, lick responses were neither rewarded nor punished. Reward probabilities during this stage were chosen from the set {0.1, 0.9} and reversed every 20–35 trials. During this period of training only, water was occasionally manually delivered to encourage learning of the response window and appropriate switching behavior. During this second stage of training, we introduced a 100-ms delay between choice and outcome. This delay was gradually increased to 300 ms for electrophysiological experiments and 200 ms for silencing and axonal calcium recordings. If a directional lick bias was observed in one session, the lick spouts were moved horizontally 50–300 μm in the opposite direction prior to the following session.

After the 3.0 or 3.1 s trial duration, inter-trial intervals (the times between consecutive cue onsets) were generated as draws from an exponential distribution with a rate parameter of 0.3 and a maximum of 20 s (and a minimum of 0.1 s in silencing experiments). This distribution results in a flat hazard rate for inter-trial intervals such that the probability of the next trial did not increase over the duration of the inter-trial interval. Inter-trial intervals were 4.83 s on average (range 1.1–20 s, including a no-lick window; see below). As in our previous studies, mice made a leftward or rightward choice in greater than 99% of trials (Bari et al., 2019; Grossman et al., 2022). Mice completed, on average, 361 trials over 51 min per session (range 153–602 trials; Figure A1).

In the final stage of the task, the reward probabilities assigned to each lick spout were drawn pseu-dorandomly from the set {0.1, 0.5,0.9}. The probabilities were assigned to each spout individually with block lengths drawn from a uniform distribution of 20–35 trials. To stagger the blocks of probability assignment for each spout, the block length for one spout in the first block of each session was drawn from a uniform distribution of 6–21 trials. For each spout, probability assignments could not be repeated across consecutive blocks. To maintain task engagement, reward probabilities of 0.1 could not be simultaneously assigned to both spouts. If one spout was assigned a reward probability greater than or equal to the reward probability of the other spout for 3 consecutive blocks, the probability of that spout was set to 0.1 to encourage switching behavior and limit the creation of a direction bias. If a mouse perseverated on a spout with reward probability of 0.1 for 4 consecutive trials, 4 trials were added to the length of both blocks. This prevented mice from choosing one spout until the reward probability became high again.

To minimize spontaneous licking, we enforced a 1 s no-lick window prior to tone onset. Licks within this window were punished with a new randomly-generated inter-trial interval, followed by a 1 s no-lick window. Implementing this window significantly reduced spontaneous licking throughout the entirety of behavioral experiments.

### Electrophysiology

We recorded extracellular signals from neurons at 32 kHz using a Digital Lynx 4SX (Neuralynx, Inc.). The recording system was connected to 8 implanted tetrodes (32 channels, nichrome wire, PX000004, Sandvik) fed through guide tubes that could be advanced with the turn of a screw on a custom, 3D-printed microdrive. The impedances of each wire in the tetrodes were reduced to 180–250 kΩ by gold plating. The tetrodes were adhered to a 200 *μ*m optic fiber used for optogenetic identification. After each recording session, the tetrode-optic-fiber bundle was driven down 30–75 *μ*m. Channels were bandpass-filtered between 0.3–6 kHz. The bandpass-filterd signal, *x*, was thresholded at 3*σ_n_* where 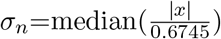 (Quiroga et al., 2004). Detected peaks were sorted into individual unit clusters offline (Spikesort 3D, Neuralynx Inc.) using peak waveform amplitude, minimum waveform trough, and waveform prinipal component analysis. We used two metrics of isolation quality as inclusion criteria: L-ratio (< 0.05) (Schmitzer-Torbert et al., 2005) and fraction of interspike interval violations (< 0.1% interspike intervals < 2 ms).

Individual neurons were determined to be identified LC-NE neurons if they responded to brief pulses (10 or 20 ms) of laser stimulation (473 nm wavelength) with short latency (< 10 ms) and high probability (> 0.8) of response to at least two pulses across trains of stimulation (10 trains of 10 pulses delivered at 5 or 10 Hz). In addition to observing responses to light stimuli, LC targeting was confirmed by performing electrolytic lesions of the tissue (25 s of 20 μA direct current across two wires of the same tetrode) and examining the tissue after perfusion.

### Viral injections

To express ChR2, GtACR2, EYFP, or GCaMP6s in LC-NE neurons, we pressure-injected 600 nl at three sites within LC of rAAV5-EF1a-DIO-hChR2(H134R)-EYFP (3 × 10^13^ GC ml^−1^), pAAV-hSyn1-SIO-stGtACR2-FusionRed (1.8 × 10^13^ GC ml^−1^, further diluted by a factor of ten), rAAV5-EF1a-DIO-eYFP (4.4 × 10^12^ GC ml^−1^, UNC viral vector core), or AAV-hSyn-FLEX-axon-GCaMP6s (1.0 × 10^13^ GC ml^−1^) into the LC of *Dbh*-Cre mice at a rate of 1 nl s^−1^ (MMO-220A, Narishige). pAAV-EF1a-double floxed-hChR2(H134R)-EYFP-WPRE-HGHpA was a gift from Karl Deisseroth (Addgene viral prep 20298-AAV5; RRID:Addgene_20298). pAAV-hSyn1-SIO-stGtACR2-FusionRed was a gift from Ofer Yizhar (Addgene viral prep 105677-AAV1; RRID:Addgene_105677) (Mahn et al., 2018). pAAV-hSynapsin1-FLEx-axon-GCaMP6s was a gift from Lin Tian (Addgene viral prep 112010-AAV5; RRID:Addgene_112010) (Broussard et al., 2018).

We made three injections of 200 nl at the following coordinates: 0.28–0.35 mm anterior to the junction of the inferior colliculus and cerebellum, 0.85–0.90 mm lateral from the midline, and {2.95, 3.15, 3.35} mm ventral from the pial surface. Before the first injection, the pipette was left at the most ventral coordinate for 5 min. Before each injection, the pipette was withdrawn 50 *μ*m and left in place for 5 min after injection. The craniotomy after a GtACR2, EYFP, or GCaMP6 injection was covered with silicone elastomer (Kwik-Cast, WPI) and dental cement. For electrophysiology experiments with rAAV5-EF1a-DIO-hChR2(H134R)-EYFP injections, the microdrive was implanted through the same craniotomy. **Inactivation of LC-NE neurons.** After training mice, four mice were injected with GtACR2 and 3 mice were injected with EYFP as a control. Optic fibers (200 μm diameter, 0.39 NA) were implanted bilaterally over LC at the same anterior-posterior and medial-lateral coordinates used for electrophysiology, using a custom 3D-printed guide holder.

### GCaMP6s measurements

To quantify calcium dynamics of LC-NE axons in prelimbic cortex, we injected AAV5-hSyn-FLEX-axon-GCaMP6s bilaterally into LC of *Dbh*-Cre mice. We implanted 200-*μ*m-diameter optic fibers (0.39 NA) above right LC (using the same coordinates as used for inactivation experiments) and above right prelimbic cortex (2.0 mm anterior to bregma, 0.5 mm lateral, 1.2 mm ventral from the pial surface). We measured GCaMP6s dynamics using 20 *μ*W irradiance of 488-nm (to excite GCaMP6s) and 405-nm (as an isosbestic control) light (Neurophotometrics). Signal was sampled at 80 Hz, alternative sampling from 405 nm and 488 nm channels.

### Data analysis

All analyses were performed with MATLAB (Mathworks), R, Python, and Stan. All data are presented as mean ± SEM unless reported otherwise. All statistical tests were two-sided. For all analyses, no-go cues were ignored and treated as part of the inter-trial interval.

### Data analysis: descriptive models of behavior

We fit logistic regression models to predict choice as a function of outcome history for each mouse using the model

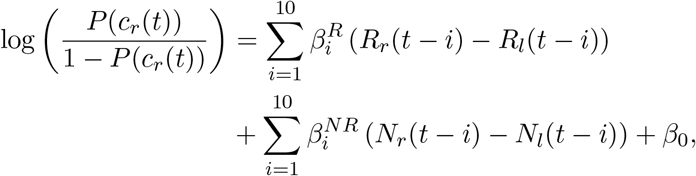

where *c_r_*(*t*) = 1 for a right choice and 0 for a left choice, *R* = 1 for a rewarded choice and 0 for an unrewarded choice, and *N* = 1 for an unrewarded choice and 0 for a rewarded choice.

### Data analysis: generative models of behavior

We applied a family of RL models of behavior used previously (Bari et al., 2019; Hattori et al., 2019; Grossman et al., 2022). This RL model estimates action values (*Q_l_*(*t*) and *Q_r_*(*t*)) on each trial to generate choices. Choices are described by a random variable, *c*(*t*), corresponding to left or right choice, *c*(*t*) ∈ {*l, r*}. The value of a choice is updated as a function of the RPE, and the rate at which this learning occurs is controlled by the learning rate parameter *α*. To account for asymmetric learning from rewards and no rewards, we used separate learning rates for each outcome. For example, if the left spout was chosen, then

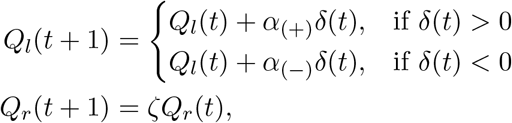

where *δ*(*t*) = *R*(*t*) – *Q_l_*(*t*) and *ζ* represents the forgetting rate parameter. The forgetting rate captures the increasing uncertainty about the value of the unchosen spout.

The *Q*-values are used to generate choice probabilities through a softmax decision function:

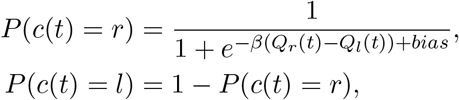

where *β*, the “inverse temperature” parameter, controls the steepness of the sigmoidal choice function. In other words, *β* controls the stochasticity of choice. This model was used for experiments with electrophysiology and GCaMP6 measurements.

For experiments with LC-NE silencing, we modified the model in two ways. First, we used a single learning parameter (*α*) to better isolate the effects of neuronal inhibition on RPE-driven behavior. Second, we added a choice auto-correlation parameter *κ* to account for the asymmetric behavior following different outcomes that was captured by separate learning rates in the previous model. Thus, for leftward choices, this model updates values according to the following equations:

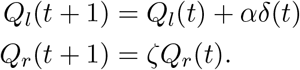

The choice function is specified as

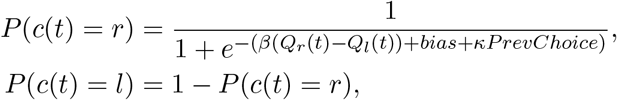

where *PrevChoice* = 1 if *c*(*t*–1) = *r* and *PrevChoice* = –1 if *c*(*t*–1) = *l*. *κ* controls choice autocorrelation, describing the trend of repeating the same choice as the one on the previous trial.

To compare change in choice dynamics against RPE, we estimated the change of choice likelihood in consecutive trials. This quantity, defined as 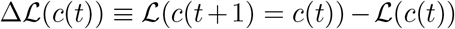, determines how future behavior is altered following RPE. 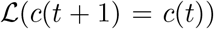 was estimated from the sequence of mouse choices by calculating the probability of choosing *c*(*t*) on trial *t* + 1 relative to trial *t*. 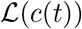 was the likelihood function estimated from the *Q*-learning model. Change of likelihood was centered for each session to account for choice autocorrelations.

To quantify the effects of LC-NE silencing, we developed two alternative models that yield different explanations for behavioral effects. The first model proposes that LC-NE silencing drives changes in RPE (Figures 4h and A4f). The second model proposes that LC-NE silencing drives changes in choice stochasticity (Figure A4g). In the first model, we added a term to *δ* (Δ_*δ*_) on trials with light delivery. In the second model, we multiplied *β* by a factor (Δ_*β*_) when light was delivered on the previous trial.

### Data analysis: model fitting

We fit and assessed models using MATLAB and the probabilistic programming language, Stan (https://mc-stan.org/) with the MATLAB interface, MatlabStan (https://mc-stan.org/users/interfaces/matlab-stan) and the GPU optimization option (Nvidia GeForce RTX 2080 Super). Stan was used to construct hierarchical models with mouse-level hyperparameters to govern session-level parameters. This hierarchical construction uses partial pooling to mitigate overfitting to noise in individual sessions (often seen in the point estimates for session-level parameters that result from other methods of estimation) without ignoring meaningful session-to-session variability. For each session, each parameter in the model (for example, *α*) was modeled as a draw from a mouse-level distribution with mean *μ* and variance *σ*. Models were fit using noninformative (uniform distribution) priors for mouse-level hyperparameters:

| Parameter | Prior distribution |
| --- | --- |
| $\alpha_{(-)}$ | $[0, 1]$ |
| $\alpha_{(+)}$ | $[0, 1]$ |
| $\alpha$ | $[0, 1]$ |
| $\zeta$ | $[0, 1]$ |
| $\beta$ | $[0, 10]$ |
| $\kappa$ | $[0, 3]$ |
| bias | $\mathcal{N}(0, 1)$ |
| $\Delta_\delta$ | $[-1, 1]$ |
| $\Delta_\beta$ | $\exp(\mathcal{N}(0, 1))$ |

Weakly informative priors were used for session-level parameters. The parameters for modeling effects of LC-NE inhibition (Δ_*δ*_, Δ_*β*_) were similarly defined with both animal-level and session-level parameters. Priors were set in [–1, 1] for Δ_*δ*_ and [0, +∞] for Δ_*β*_ to allow the model to capture potentially bidirectional effects of silencing and quantify the effects of LC-NE silencing (i.e., whether the posterior distribution of Δ_*δ*_ differs from zero and whether the posterior of Δ_*β*_ differs from 1). Mouse-level hyperparameters were chosen to achieve model convergence under the assumption that individual mice behave similarly across days. The parameters were sampled in an unconstrained space and transformed into bounded values (for those parameters that were bounded) by a standard normal inverse cumulative density function. Stan uses full Bayesian statistical inference to generate posterior distributions of parameter estimates using Hamiltonian Markov chain Monte Carlo sampling (Carpenter et al., 2017). The default no-U-turn sampler was used. The Metropolis acceptance rate was set to 0.85–0.9 to force smaller step sizes and improve sampler efficiency. The models were fit with 10,000 iterations and 5,000 warmup draws run on each of 8 chains in parallel. Default configuration settings were used otherwise.

In Figures 4i, A4e, and A4h, we included an experiment-level prior (uniform distribution on [–1,1] for Δ_*δ*_ and 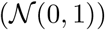 for Δ_*β*_) and we did not include session-level variation of parameters within-mouse. We included an experiment-level prior here to account for the fact that the same experimental manipulation (light delivery) was applied in each mouse.

### Data analysis: extracting model parameters and variables, behavior simulation

For extracting model variables (like RPE), we took at least 2,000 draws from the Hamiltonian Markov chain Monte Carlo samples of session-level parameters, ran the model agent through the task with the actual choices and outcomes, and averaged each model variable across runs. For comparisons of individual parameters across models, we estimated *maximum a posteriori* parameter values by approximating the mode of the distribution: binning the values in 50 bins and taking the mean value of the most populated bin. For simulations of effects of LC-NE silencing, we took on average 1000 draws (varied in proportion with the number of sessions collected from each mouse) from the Hamiltonian Markov chain Monte Carlo samples of mouse-level parameters and simulated behavior and outcomes in one session per sample.

### Data analysis: linear regression models of firing rates

To characterize neuronal responses to choice outcomes, we regressed firing rates on outcome (*R*), reward expectation (*Q_c_*) and choice (*c*(*t*)) using the MATLAB function “fitlm”. We quantified regressions using both t-statistics of regressors (Figure 3b) and the angle in Cartesian coordinates of the estimates of coefficients for *R* and *Q_c_* for each neuron. The latter was plotted as a polar histogram (Figure 3c).

### Data analysis: GCaMP6s dynamics

Raw fluorescence was low-pass filtered at 10 Hz. Both channels were z-scored and a linear model was fit to the z-scored signals using the 488-nm channel as the dependent variable, the 405-nm channel as the regressor, together with an intercept. An estimate of the baseline fluorescence was subtracted from the model residuals. To estimate the baseline, we calculated the largest tenth percentile of the residuals in a 15-s sliding window.

### Data analysis: estimates of GCaMP6s dynamics from firing rates

To make predictions about GCaMP6s dynamics from measured firing rates (Figure 3d), we used a spike-to-fluorescence transfer function (Wei et al., 2020). We first generated a latent variable by convolving a causal filter with firing rates from each neuron in each population of LC-NE neurons. We then passed the latent variable through an activation function. This yielded a predicted GCaMP6s trace for each neuron. We then z-scored and averaged type I and type II neurons separately (Figure 5c).

### Data analysis: spike count correlations

To estimate firing rate correlations among simultaneously recorded LC-NE neurons, we counted spikes in a 2-s pre-cue window and calculated correlation coefficients among simultaneously recorded type I or type II neurons. For visualization (Figure A2d), we smoothed trial-by-trial baseline firing rates using a boxcar filter in 10-trial bins.

### Histology

After experiments were completed, mice were euthanized with an overdose of isoflurane, exsanguinated with saline, and perfused with 4% paraformaldehyde. The brains were cut in 100-μm-thick coronal sections and mounted on glass slides. We validated expression of rAAV5-EF1a-DIO-hChR2(H134R)-EYFP, pAAV-hSyn-SIO-GtACR2-FusionRed, or pAAV-hSyn-DIO-EYFP with epifluorescence images of LC (Zeiss Axio Zoom.V16). In electrophysiological experiments, we confirmed targeting of the optic-fiber-tetrode bundle to LC by colocalization of the electrolytic lesion with immunostaining against tyrosine hydroxylase (rabbit anti-TH, ab152, Millipore, followed by donkey anti-rabbit Alexa 488, Life Technologies, or goat anti-rabbit Alexa 594, Life Technologies). To visualize GCaMP6s expression in axons in PrL, we used chicken anti-GFP (ab13970, Abcam) with goat anti-chicken Alexa 488 (Life Technologies).

---

**Figure A1.**
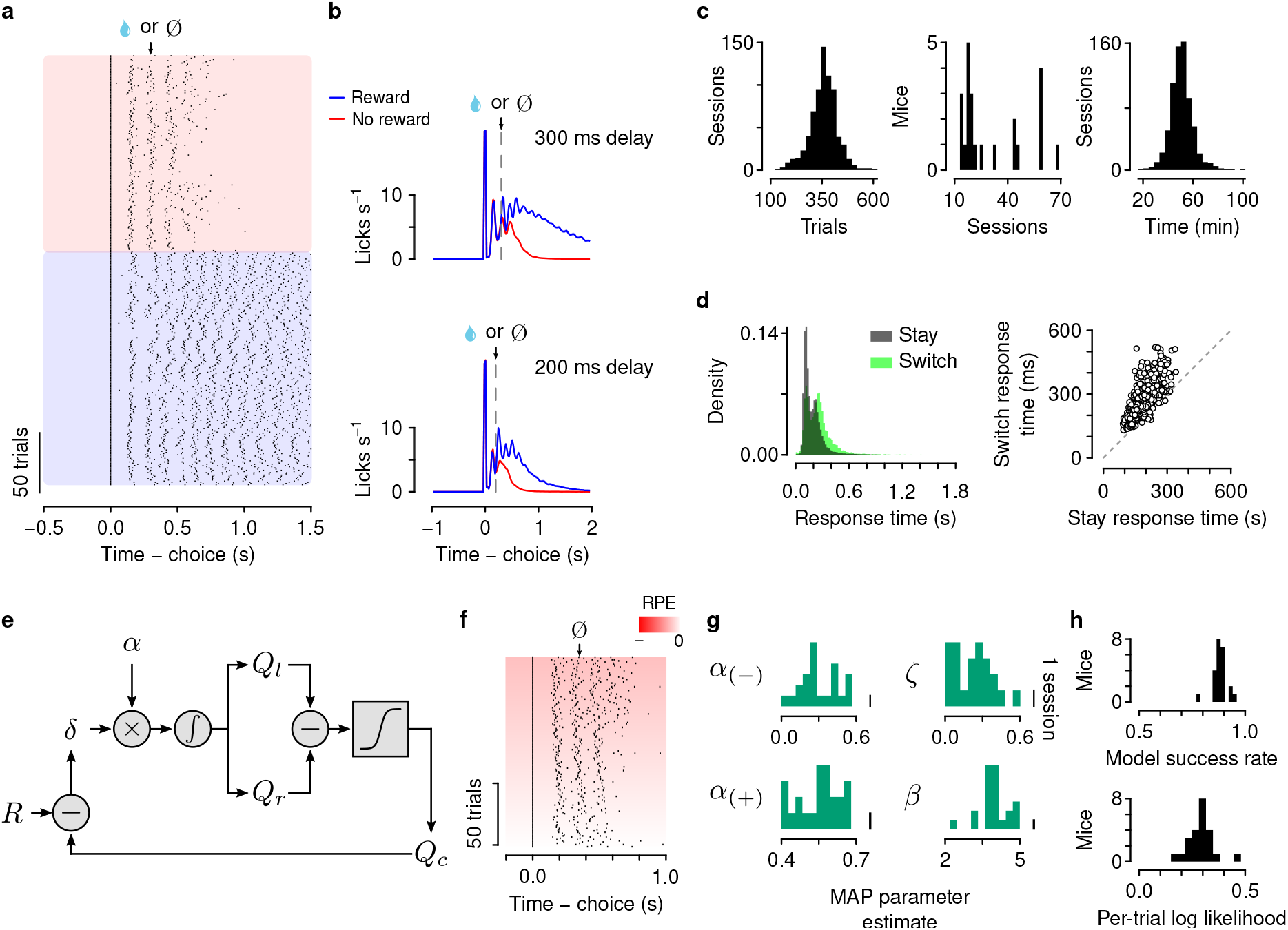
Behavioral performance and *Q*-learning model fitting. (a) Example lick times on rewarded (blue) and unrewarded (red) trials. (b) Mean ± SEM lick rates on rewarded and unrewarded trials with a 300-ms delay (top) or 200-ms delay (bottom) between choice and outcome. (c) Number of trials per session (left), sessions per mouse (middle), and time per session (right). (d) Response times on trials in which mice switched choice relative to the previous one were slower than when they repeated the same choice (607 sessions, paired t-test, *t*_606_ = 38.53, *p* = 4.4 × 10^−165^). (e) Reinforcement-learning model. Key variables include reward (*R*), chosen value (*Q_c_*), reward prediction error (RPE, *δ*), and values of left (*Q_l_*) and right (*Q_r_*) actions. *Relative value* (*Q_r_* – *Q_l_*) is used to make choices through a softmax decision function. The predicted value of a choice (*Q_c_*) is compared to reward (*R*) to generate a *reward prediction error* (*δ*, RPE). (f) Example lick raster from all unrewarded trials from one session, showing more licks following more negative RPE. (g) *Maximum a posteriori* (MAP) estimates of model parameters across mice. (h) Model success rate (probability that model choices equal mouse choices) and per-trial negative log likelihood across mice.

**Figure A2.**
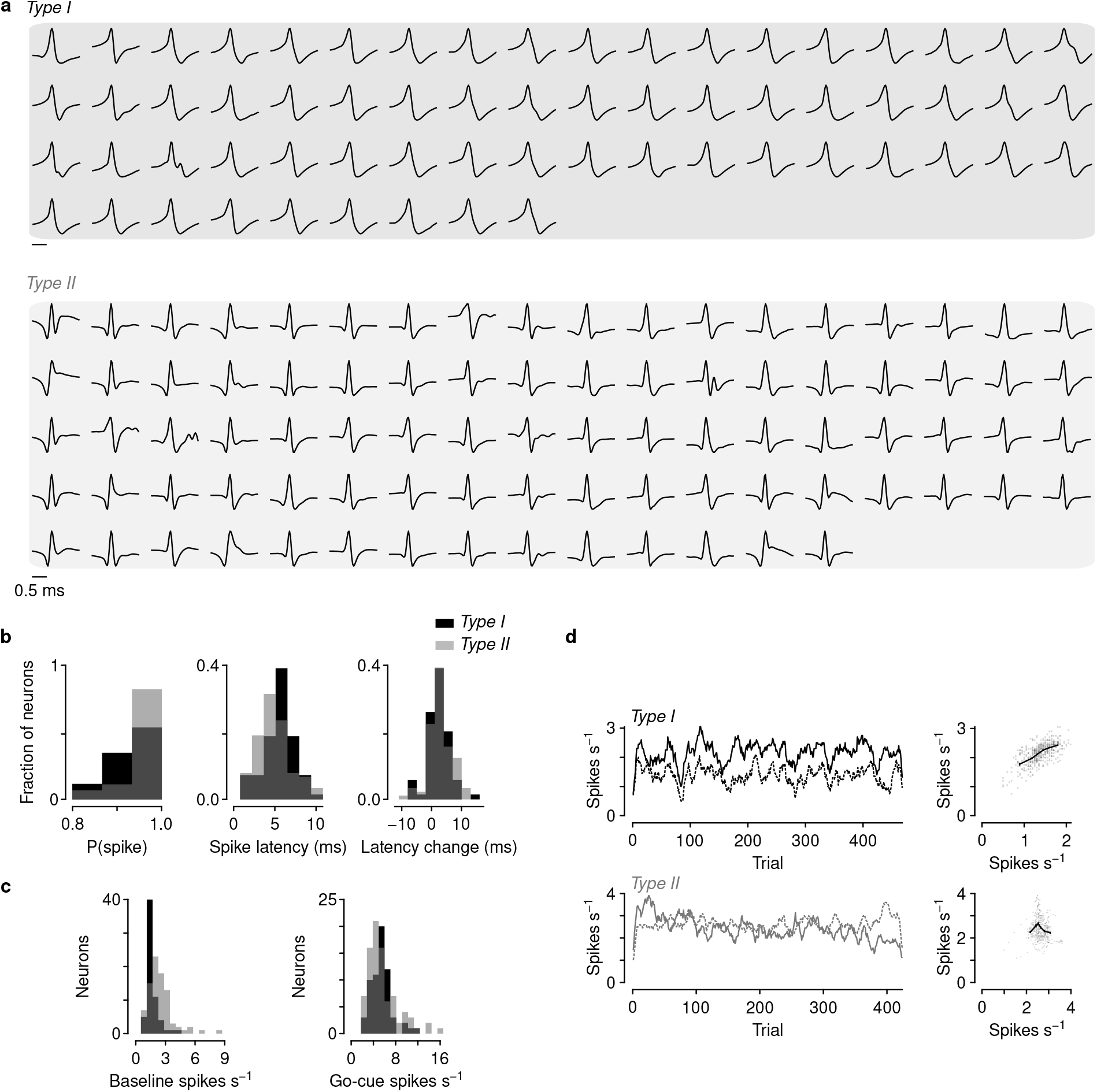
Two types of LC-NE neurons. (a) Mean action potential waveforms from all type I and type II neurons. (b) Probability of firing in response to light stimuli (left), spike latency (middle), and increase in latency with successive pulses in a train (right) among identified LC-NE neurons. (c) Baseline firing rates were significantly higher among type II than type I neurons (t-test, *t*_147_ = 4.32, *p* = 2.9 × 10^−5^). Firing rates following go cues were not significantly different (t-test, *t*_147_ = 0.86, *p* = 0.39). (d) Firing rates in the 2 s before go cues from example simultaneously recorded pairs of type I and type II neurons (10-trial boxcar filter). Scale bar, 1 z-score. Note the stronger correlations between type I (correlation coefficient, 0.68) versus type II neurons (correlation coefficient, –0.024).

**Figure A3.**
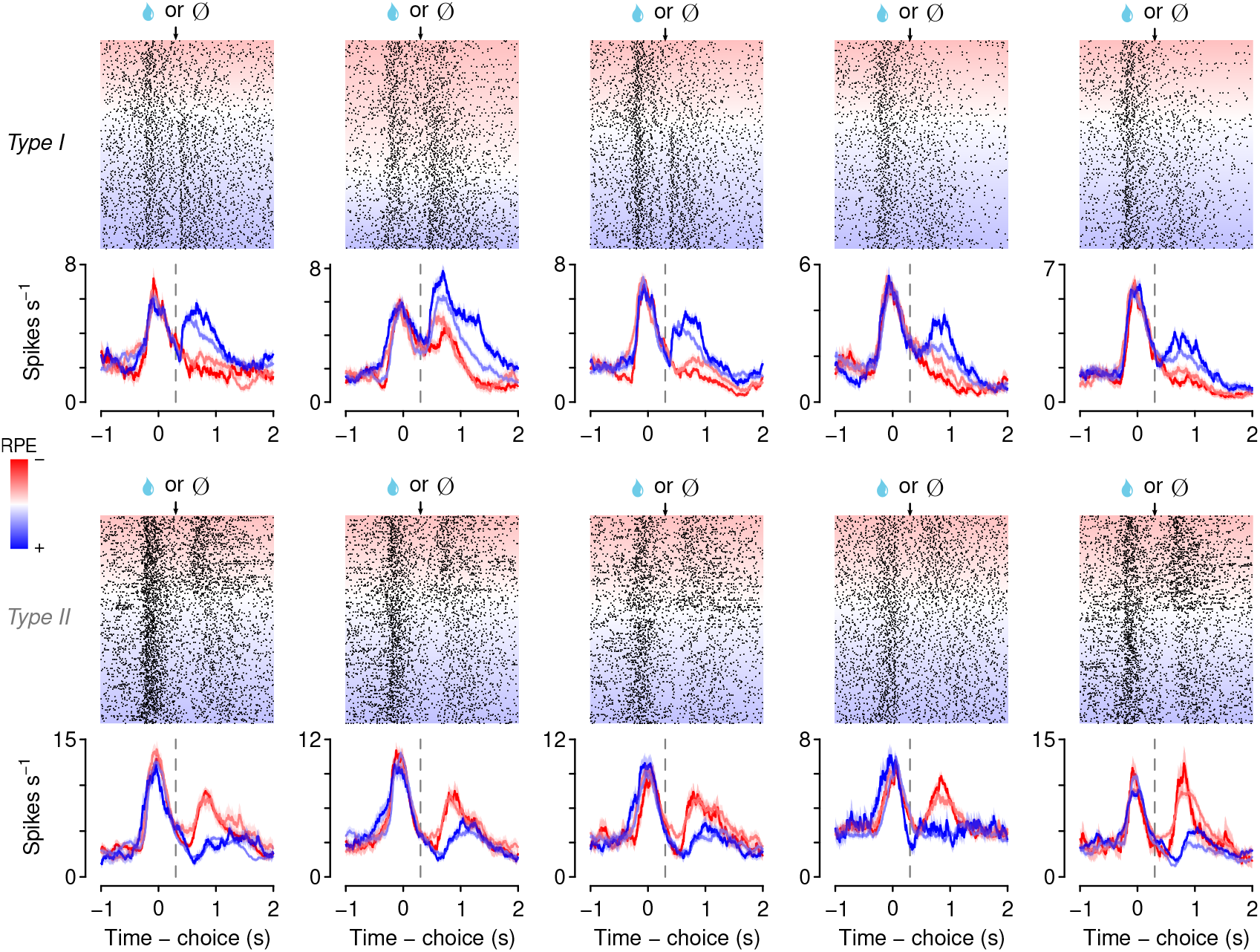
Example neurons of each type. (a) Type I neurons correlated with RPE. Type II neurons were excited by no reward. Examples from Figure 3 are reproduced here for comparison.

**Figure A4.**
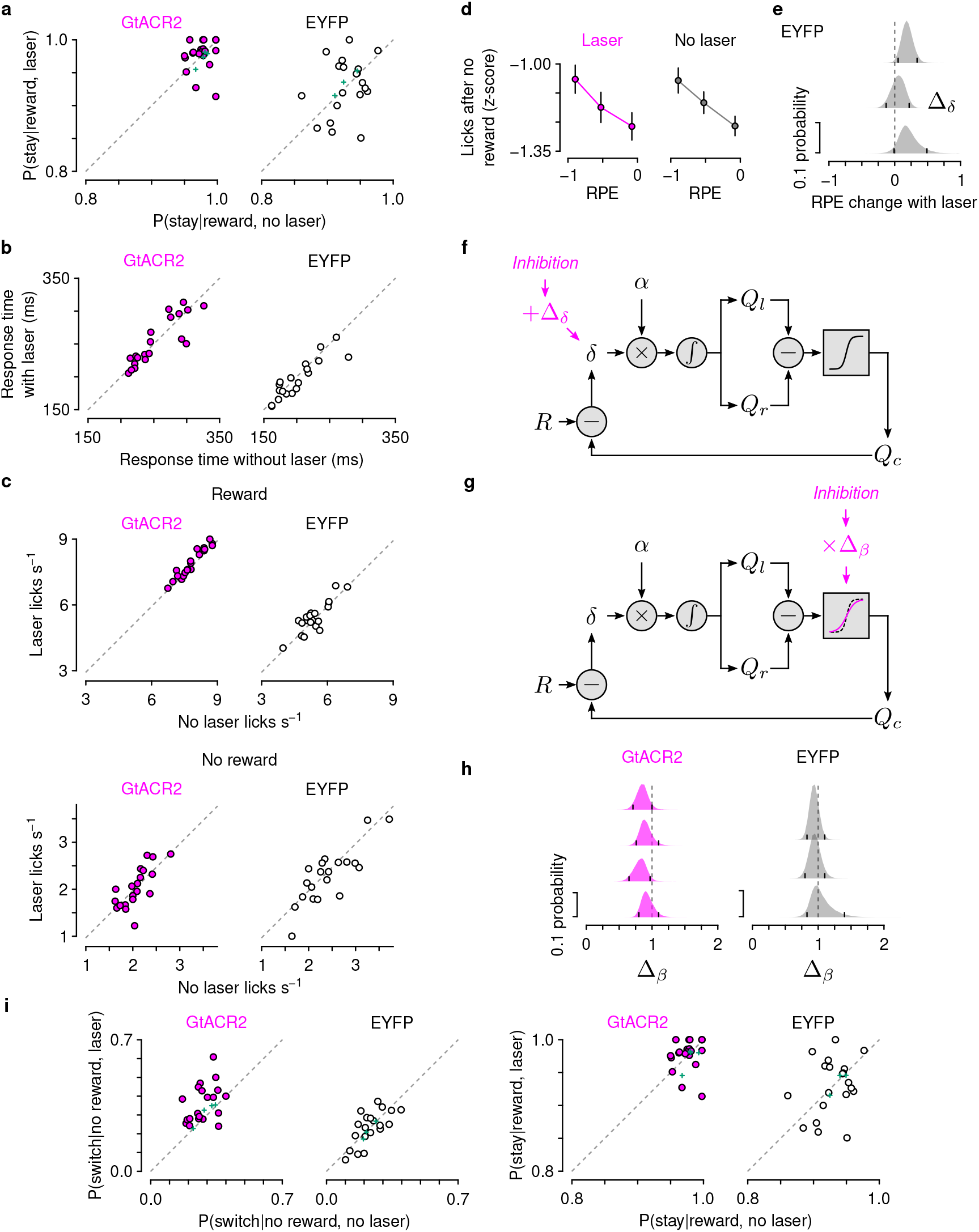
LC-NE inactivation did not disrupt ongoing movements or choice stochasticity. (a) No change in *P*(stay|reward) with silencing (signed rank tests, GtACR2, *p* = 0.45; EYFP, *p* = 0.88). Green: predictions from the model with an additive change in RPE. (b) No change in response time with silencing (Wilcoxon signed rank tests: GtACR2, 0.79; EYFP, *p* = 0.18). (c) No change in number of licks on rewarded or unrewarded trials (signed-rank tests: GtACR2, reward, *p* = 0.093, no reward, *p* = 0.74; EYFP, reward, *p* = 0.65, no reward, *p* = 0.13). (d) The increase in lick number with increased negative RPE following no reward remained with silencing (linear regression coefficients: laser, –0.23 ± 0.11, *t* = –2.2, *p* = 0.030; no laser, –0.38 ± 0.11, *t* = –5.3, *p* = 1.4 × 10^−7^). (e) Light delivery did not negatively shift RPE in fluorophore-control mice. Ticks correspond to 95% certainty intervals. (f) A model in which silencing is hypothesized to change RPE (Figure 4h). (g) A model in which silencing is hypothesized to change choice randomness (by affecting the choice function) did not show a strong change in choice stochasticity with LC-NE silencing (h). Ticks correspond to 95% certainty intervals. (i) *P*(switch|no reward) and *P*(stay|reward) for trials with and without laser for each group of mice. Mouse data are reproduced from Figures 4f and A4a. Green: predictions from model in (g).

**Figure A5.**
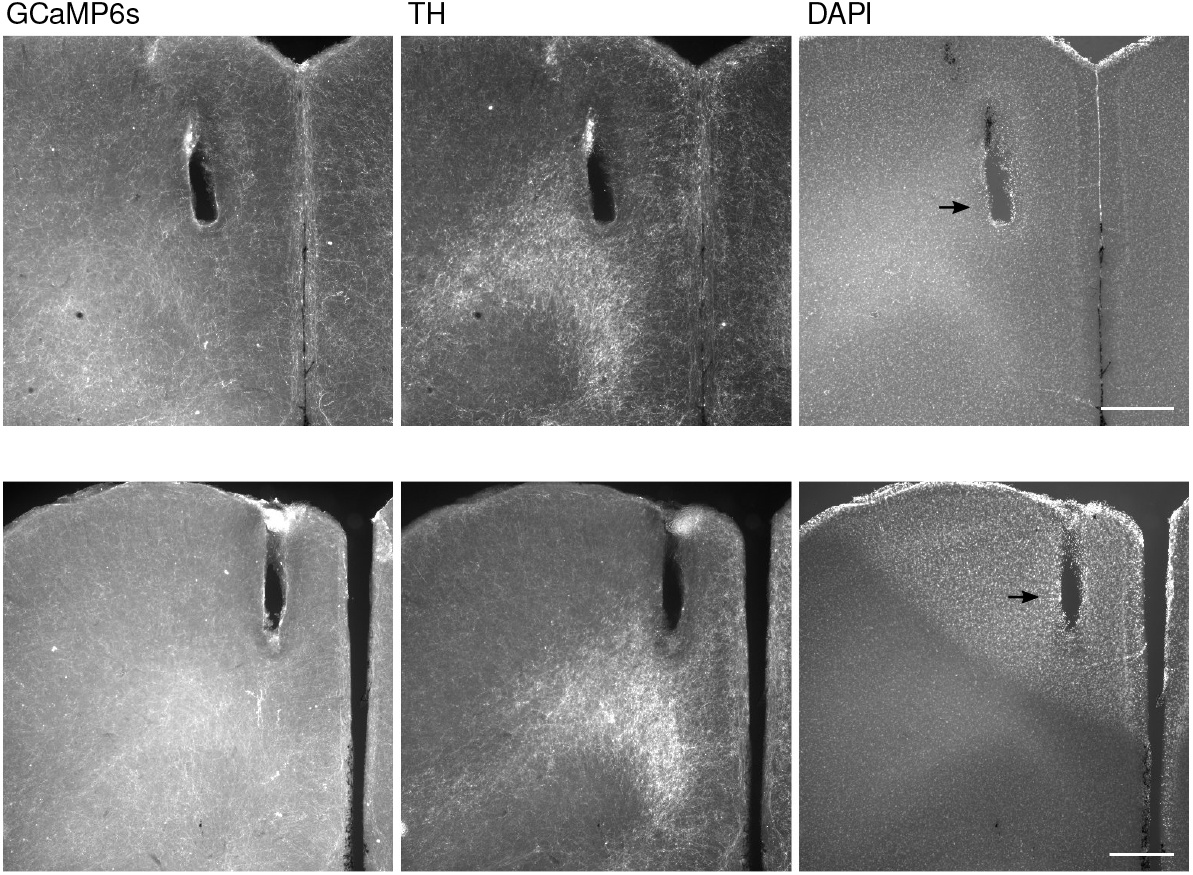
Expression of GCaMP6s in LC-NE axons in PrL, together with TH immunostaining and DAPI to label nuclei. Arrowheads: fiber optic track. Scale bars: 0.5 mm.

